# Conservation of sporulation genes and a transmembrane-containing Spo0B variant in Paenibacillus

**DOI:** 10.1101/2025.08.24.672004

**Authors:** Isabella N. Lin, Cassidy R. Prince, Heather A. Feaga

## Abstract

Sporulation is a strategy employed by many bacteria to survive harsh environmental conditions. The genus *Paenibacillus* includes spore-forming species notorious for spoiling pasteurized dairy products and causing fatal infections in honeybee larvae, leading to colony collapse. Here, we present a comprehensive survey of sporulation genes across 1460 high-quality *Paenibacillus* genomes. We find that all members of the sporulation-initiating phosphorelay are well-conserved, but that the Spo0B phosphotransferase contains a transmembrane domain that is unique to this genus. The transmembrane-domain-containing variant of Spo0B is present in 92% of surveyed *Paenibacillus* genomes. Consistent with this high level of conservation, we find that for *Paenibacillus polymyxa* Spo0B, the transmembrane domain is important for interaction with its phosphorelay partners Spo0A and Spo0F. Moreover, we find that Spo0B exhibits low sequence identity across Bacilli when compared to other members of the phosphorelay. Altogether, this work highlights the potential for diversity even within the highly conserved phosphorelay that initiates sporulation in Bacilli.

**Importance:** Spores are the most durable life-form, and the sporulation process serves as a paradigm of cellular development and differentiation. Sporulation is well-characterized in the model organism *Bacillus subtilis*, but we lack information about non-model spore-formers. The genus *Paenibacillus* includes spore-formers that negatively impact farming and food industries. Here, we present the first comprehensive search for sporulation genes in *Paenibacillus* and show that a unique transmembrane-domain-containing Spo0B is widespread throughout this genus.

## Introduction

Endospore formation is a complex developmental process limited to certain classes of Firmicutes (1, 2). Through sporulation, vegetative bacteria undergo developmental and metabolic changes to produce highly resistant, dormant spores that are capable of withstanding adverse environmental conditions (3). Spore formation involves a series of morphological stages that is typically triggered by low nutrient density. A vegetative cell asymmetrically divides into a mother cell and forespore, after which the forespore is fully engulfed and released into the mother cell cytoplasm (4–7). The forespore is encased in two protective shells, the cortex and the coat (8–10), and the mature spore is released via lysis of the mother cell (9). Upon sensing germinants, typically nutrients, the spore will germinate and outgrow into a vegetative cell (11, 12).

Endospore formation is well-characterized in *Bacillus subtilis,* where it affects the expression of over 500 genes (13, 14). Sporulation-associated genes continue to be identified and characterized (15–20). A transposon screen by Meeske et al. in 2016 identified 157 genes required for the sporulation of *B. subtilis,* including 24 previously unidentified genes (21). Genes required for sporulation include kinases and phosphotransferases that initiate sporulation, sigma factors that control endospore progression, chromosome segregation, and production of the spore coat. Many of these proteins have dedicated roles in sporulation, while others, such as Spo0A, also play a role in vegetative growth (22–24).

*B. subtilis*, along with many other Bacilli and Clostridia, initiates sporulation via a phosphorelay system (25–27). Five autophosphorylating sensor histidine kinases (KinA-KinE) respond to high cell density and low nutrient availability by transferring a phosphoryl group to Spo0A via the phosphotransferases Spo0F and Spo0B (4, 27–34). Spo0A, the key transcriptional regulator of sporulation, is activated in its phosphorylated form and directly influences the expression of over 100 genes to govern entry into sporulation (35, 36). In Bacilli and Clostridia, sporulation kinases are highly conserved, as is Spo0A, which is found even in non-sporulating members of the Firmicutes (1). In contrast, many species within Clostridia lack the phosphotransferases and instead multiple kinases phosphorylate Spo0A directly (25, 37–39).

*Paenibacillus* is a spore-forming genus of bacteria encompassing over 200 species with diverse characteristics (40). *Paenibacillus* is commonly found in soil, and many species promote the growth of plants through phosphate solubilization or nitrogen fixation, such as *P. polymyxa*, *P. macerans*, and *P. elgii* (41–43). Many species also contribute to biological control by producing biocidal compounds, including *P. alvei*, *P. ehimensis*, and *P. kribbensis* (44–46). While generally non-pathogenic, *Paenibacillus* has been documented to infect humans, but is typically an opportunistic infector (47–49). *Paenibacillus* is also the etiological agent of American Foulbrood Disease, a devastating honeybee brood disease caused by *P. larvae,* for which there is currently no treatment and necessitates burning of infected hives (50, 51). Species such as *P. odorifer*, *P. amylolyticus*, and *P. lactis* are the causative spoilage agents of a variety of food products, including pasteurized and chilled products such as dairy and ready-to-eat meals (52–54). Studying the sporulation of diverse *Paenibacillus* can have impacts across many industries.

Here, we survey 1460 high-quality *Paenibacillus* genomes for known sporulation-associated protein-coding genes from *B. subtilis*. We detected 632 *B. subtilis* sporulation genes in *Paenibacillus*, with 350 of these genes conserved in ≥95% of the genomes. Specifically, we show that a Spo0B variant containing a unique N-terminal transmembrane region (Spo0B-TM) is conserved across 92% of *Paenibacillus* genomes. This Spo0B-TM variant was not functionally interchangeable with Spo0B in *B. subtilis*, but Spo0B-TM associates with its phosphorelay partners, Spo0F and Spo0A, from *P. polymyxa* and the transmembrane domain is required to detect this association. Across Spo0B orthologs in Bacilli, we found that while the phosphorylatable histidine region is strongly conserved in sequence, the rest of the protein is highly variable between species and, in the case of Spo0B from *Paenibacillus*, this sequence variation is likely consequential.

## Results

### Survey of the *Paenibacillus* sporulation pangenome identifies core sporulation genes that are shared with other spore-forming Firmicutes

To survey the conservation of known sporulation genes throughout the *Paenibacillus* genus, we downloaded all high-quality annotated *Paenibacillus* genome assemblies (n = 1460) from NCBI and performed a BLASTP search using 741 *B. subtilis* sporulation protein sequences. Genes encoding the protein sequences used in our search included genes predicted to be essential for sporulation in *B. subtilis* (21), genes classified by SubtiWiki as sporulation genes (55), and genes in the sporulation gene set defined by Galperin et al. (23). The surveyed *Paenibacillus* genomes represent over 250 unique species (Table S1), with strong representation from *P. polymyxa* (n = 117), *P. larvae* (n = 64), and *P. odorifer* (n = 37). As additional genomes were included in the search, fewer additional sporulation genes were detected, indicating that the search was performed to saturation. Thus, the number and quality of genomes surveyed were sufficient to capture an accurate, extensive representation of likely sporulation genes within the genus that are shared with *B. subtilis*.

Of the 741 *B. subtilis* sporulation genes included in our search, 632 were detected in at least one of the *Paenibacillus* genomes (Table S2), with a mean of 453 sporulation genes detected per genome. To determine which known sporulation genes may be most important for sporulation in *Paenibacillus*, we split the sporulation pangenome into core, shell, and cloud genomes. Genes found in ≥95% of the surveyed genomes made up the core, genes found in <95% but ≥15% of genomes made up the shell, and the remaining genes made up the cloud. Galperin et al. (23) constructed a list of ∼120 universally conserved sporulation genes in Firmicutes. We detected all 120 of these genes in *Paenibacillus* but found 115 of these genes in the core sporulation genome. Over half of the sporulation genes we identified were core genes (n = 350), of which 14 are essential for life in *B. subtilis* (56). Meeske et al. identified 142 genes required for sporulation in *B. subtilis,* excluding genes involved in the TCA cycle (21). We found 127 of these genes in the sporulation core for *Paenibacillus*. The key players in the phosphorelay, KinA-E, and Spo0F, Spo0B, and Spo0A are conserved in >99% of *Paenibacillus* genomes we surveyed (Fig 1, Fig 2, Table S2). The sporulation specific sigma factors, SigE, SigF, SigK, and SigG (57) are conserved in 100% of the genomes (Table S2). These results indicate that *Paenibacillus* shares a very similar sporulation pathway with *B. subtilis*.

**Figure 1.**
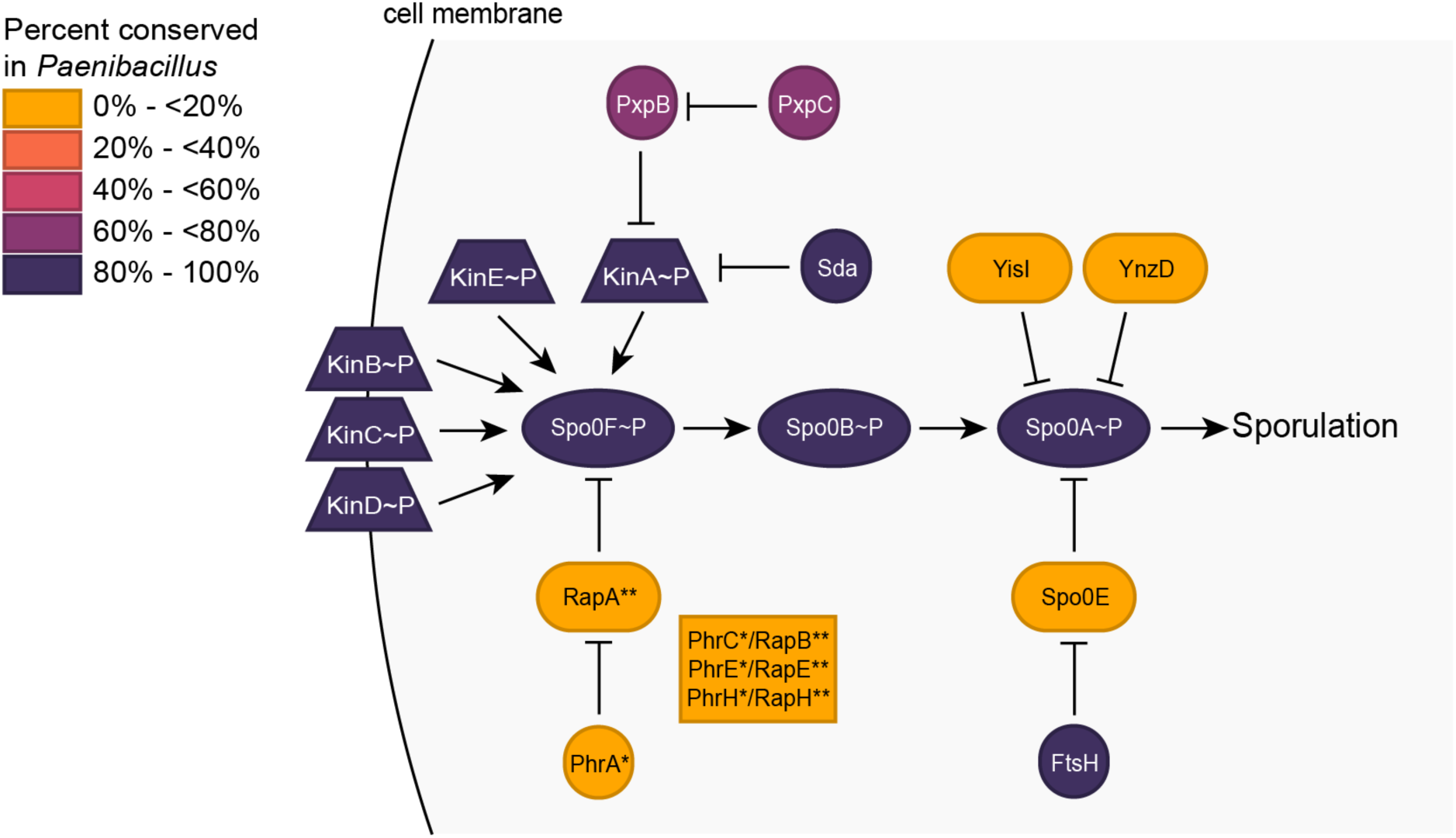
Key sporulation initiation proteins in *B. subtilis* are conserved in *Paenibacillus*. Heat map of the sporulation initiation phosphorelay of *B. subtilis*, showing the conservation of the proteins across *Paenibacillus*.

**Figure 2.**
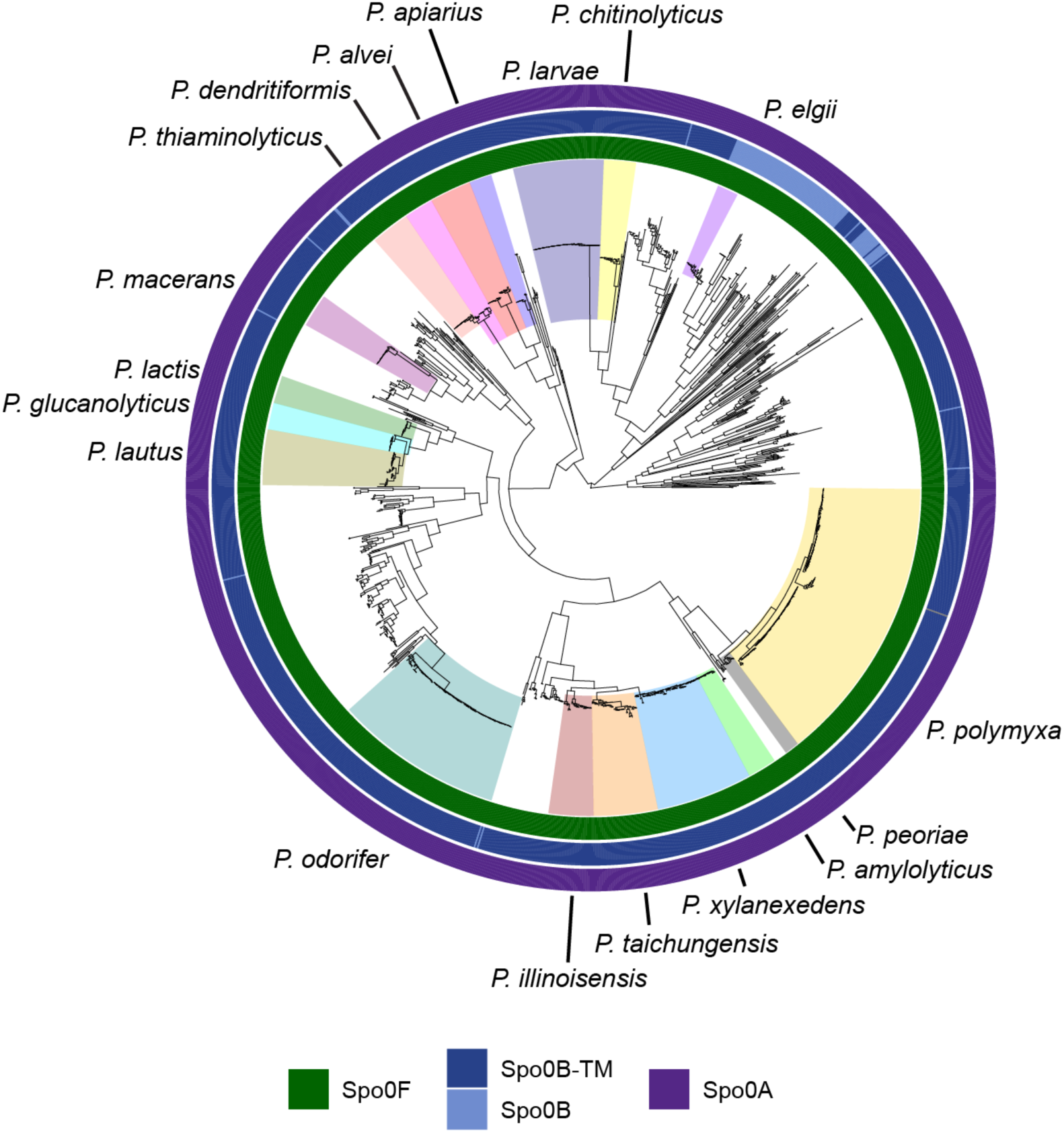
**Phosphorelay proteins are conserved in *Paenibacillus.*** A midpoint rooted maximum likelihood phylogenetic tree of *Paenibacillus* genomes built with single copy core genes. Species with at least nine available high-quality genomes are labeled. Heat maps show the presence of Spo0F, Spo0B, and Spo0A, and the transmembrane domain variant of Spo0B.

### A Spo0B-variant with an N-terminal transmembrane region is conserved across *Paenibacillus*

A previous study of sporulation histidine kinases of *P. polymyxa* examined the Spo0B sequences of six *Paenibacillus* genomes and found a unique extended N-terminal region of the protein containing two hydrophobic helices that are predicted to form a transmembrane domain (Fig 3A) (58). We will refer to Spo0B containing the transmembrane region as Spo0B-TM. Of the *Paenibacillus* genomes surveyed, 92% encoded a Spo0B variant with a predicted N-terminal transmembrane region (Fig 2). The remaining 8% of *Paenibacillus* genomes encoded a Spo0B that lacked the transmembrane domain. Species that do not contain the transmembrane region of Spo0B clade together and include *P. elgii*, *P. validus, P. mucilaginosus, P. ehimensis*, and *P. tyrfis* (Fig 2).

**Figure 3.**
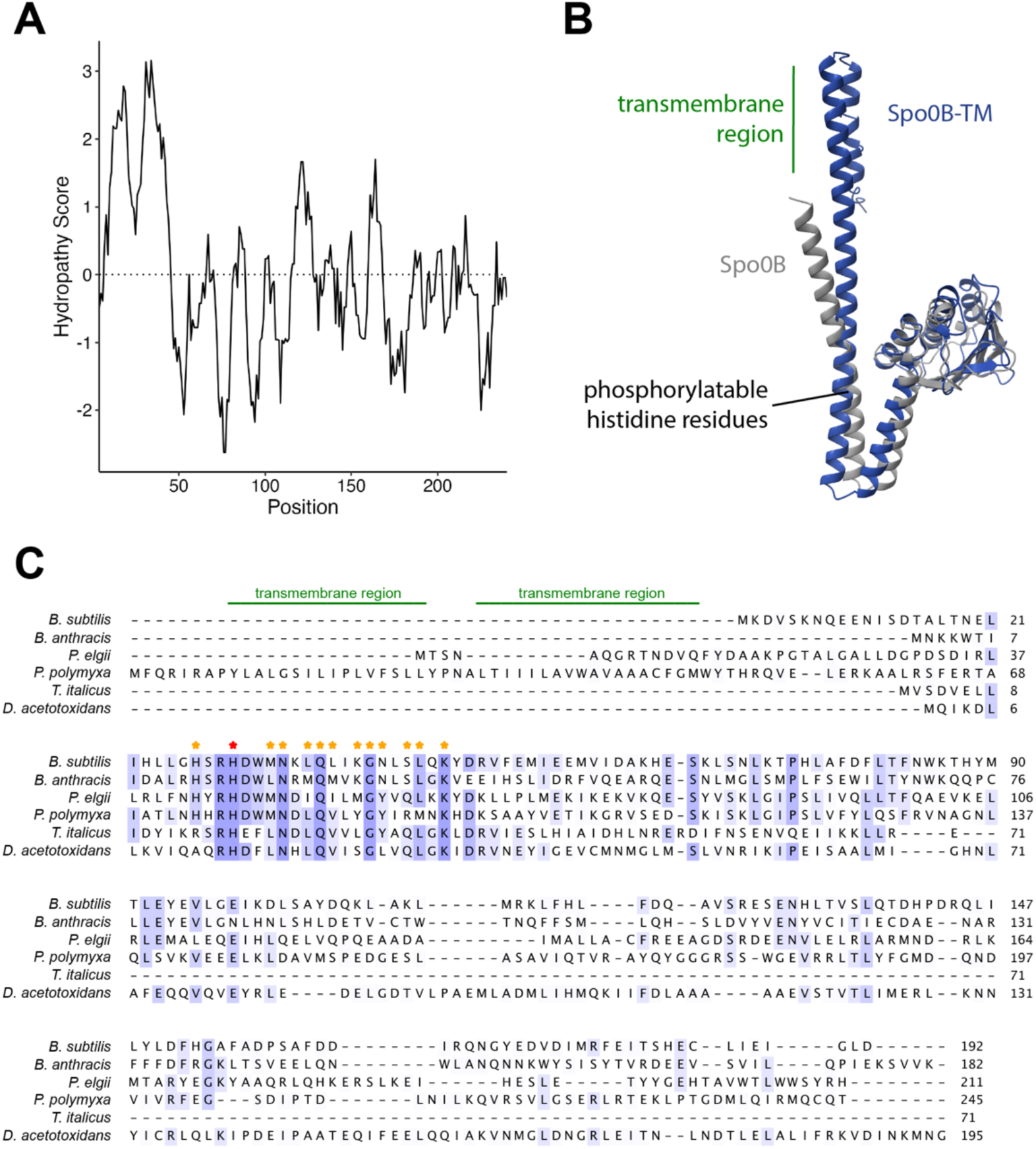
Spo0B has a unique transmembrane domain in most *Paenibacillus* genomes. **(A)** Kyte-Doolittle hydropathy plot of Spo0B-TM from *P. polymyxa.* **(B)** AlphaFold model of Spo0B from *B. subtilis* and Spo0B-TM from *P. polymyxa.* **(C)** Protein sequence alignment of Spo0B in *B. subtilis* and its orthologs across Bacilli (*Bacillus anthracis*, *Paenibacillus elgii*, *Paenibacillus polymyxa*) and Clostridia (*Thermoanaerobacter italicus, Desulfofarcimen acetotoxidans*). Amino acids are shaded based on conservation, with completely conserved residues shown in dark purple. The transmembrane region of Spo0B-TM from *P. polymyxa* is annotated. Residues in the α1 helix region of Spo0B from *B. subtilis* that are known to interact with Spo0F and Spo0A are indicated with stars. The phosphorylatable histidine residue is denoted by the red star.

Despite low percent identity at the protein sequence level (19%), Spo0B from *B. subtilis* and Spo0B-TM from *P. polymyxa* are predicted to have a high degree of structural similarity when modeled using AlphaFold (Fig 3B). Alignment of six Spo0B orthologs from Bacilli and Clostridia revealed a highly conserved region around the phosphorylatable histidine residue (Fig 3C). As expected, the residues that directly interact with Spo0F and Spo0A in this α1 helix region in *B. subtilis* (59) are completely or highly conserved (Fig 3C).

### Spo0B-TM interacts with phosphorelay proteins Spo0A and Spo0F from *P. polymyxa*

To predict if *P. polymyxa* Spo0F and Spo0A could interact with Spo0B-TM, we modeled the interactions using AlphaFold (Fig 4A). The interaction between Spo0F and Spo0B-TM had an ipTM score >0.8, indicating a confident, high-quality prediction (60). The interaction between Spo0A and Spo0B-TM had an ipTM score of 0.66, indicating the prediction was not made with high confidence. However, the interaction model of *B. subtilis* Spo0A and Spo0B, proteins known to interact, similarly yielded an ipTM score of 0.63. Therefore, the low score of the Spo0A and Spo0B-TM interaction should not discredit the modeled interaction. The location of the binding between the two protein pairs across species also appeared similar (Fig 4A).

**Figure 4.**
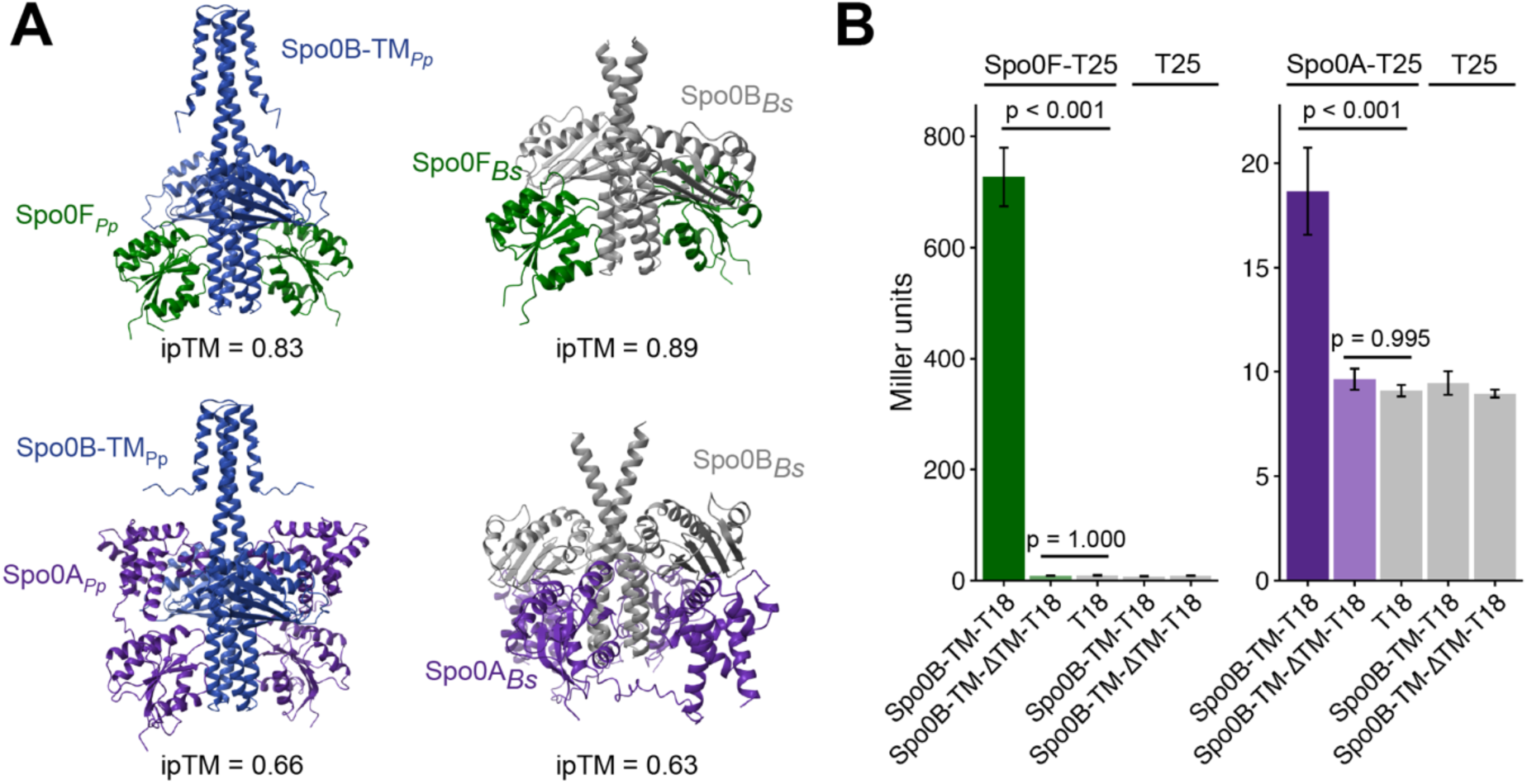
Spo0B-TM interacts with both Spo0F and Spo0A from *P. polymyxa*. **(A)** AlphaFold models of *P. polymyxa* and *B. subtilis* protein interactions. ipTM scores >0.8 represent high confidence predictions. *Pp, P. polymyxa; Bs, B. subtilis*. **(B)** Quantification of β-galactosidase activity for BACTH constructs using *P. polymyxa* proteins, representative of nine biological replicates. Error bars represent SEM. *P*-values indicate the results of a one-way ANOVA followed by Tukey’s test.

To examine the interaction of Spo0B-TM with Spo0A and Spo0F from *P. polymyxa in vivo*, we used a bacterial adenylate cyclase two-hybrid (BACTH) system in *Escherichia coli*. The BACTH system utilizes the interaction-mediated reconstitution of adenylate cyclase from *Bordetella pertussis* (61). The catalytic domain of this protein is active only when its two complementary fragments (T18 and T25) are brought together to allow for functional complementation. When interacting proteins are fused to T18 and T25, heterodimerization of the proteins results in functional complementation of the adenylate cyclase fragments. Adenylate cyclase synthesizes cAMP which induces the production of β-galactosidase in the cell. Therefore, we used β-galactosidase activity as a reporter to quantify protein-protein interactions.

We fused *P. polymyxa* Spo0F or Spo0A to the T25 fragment and *P. polymyxa* Spo0B-TM to the T18 fragment of the adenylate cyclase catalytic domain. Spo0B-TM showed strong interaction with Spo0F, with a β-galactosidase activity of 727 ± 53 Miller units (Fig 4B). The interaction of Spo0B-TM with Spo0A yielded a more modest level of β-galactosidase activity (19 ± 2 Miller units), but this activity was significantly higher than the negative controls (p < 0.001 compared to all negative controls) (Fig 4B). Interestingly, Spo0B-TM did not exhibit detectable interaction with either Spo0A or Spo0F when the transmembrane region was deleted (Spo0B-TMΔTM), with β-galactosidase activity comparable to the negative controls (Fig 4B). These results suggest that Spo0B-TM interacts with Spo0A and Spo0F of *P. polymyxa*, and that the transmembrane domain is necessary for this interaction.

### Spo0B-TM is syntenic, but not functionally interchangeable with Spo0B in *B. subtilis*

While two-component signaling proteins are typically organized in adjacent operons (62), the sporulation initiation phosphorelay encoding genes are scattered throughout the genome. In *B. subtilis*, *spo0B* is located downstream of the *rplU* and *rpmE*, genes encoding ribosomal proteins bL21 and bL31 (63, 64), and upstream of *obg*, the gene encoding the GTP-binding protein Obg (65). We examined the syntenic regions in Bacilli and Clostridia species and found the genomic arrangement of *spo0B* to be well-conserved (Fig 5A).

**Figure 5.**
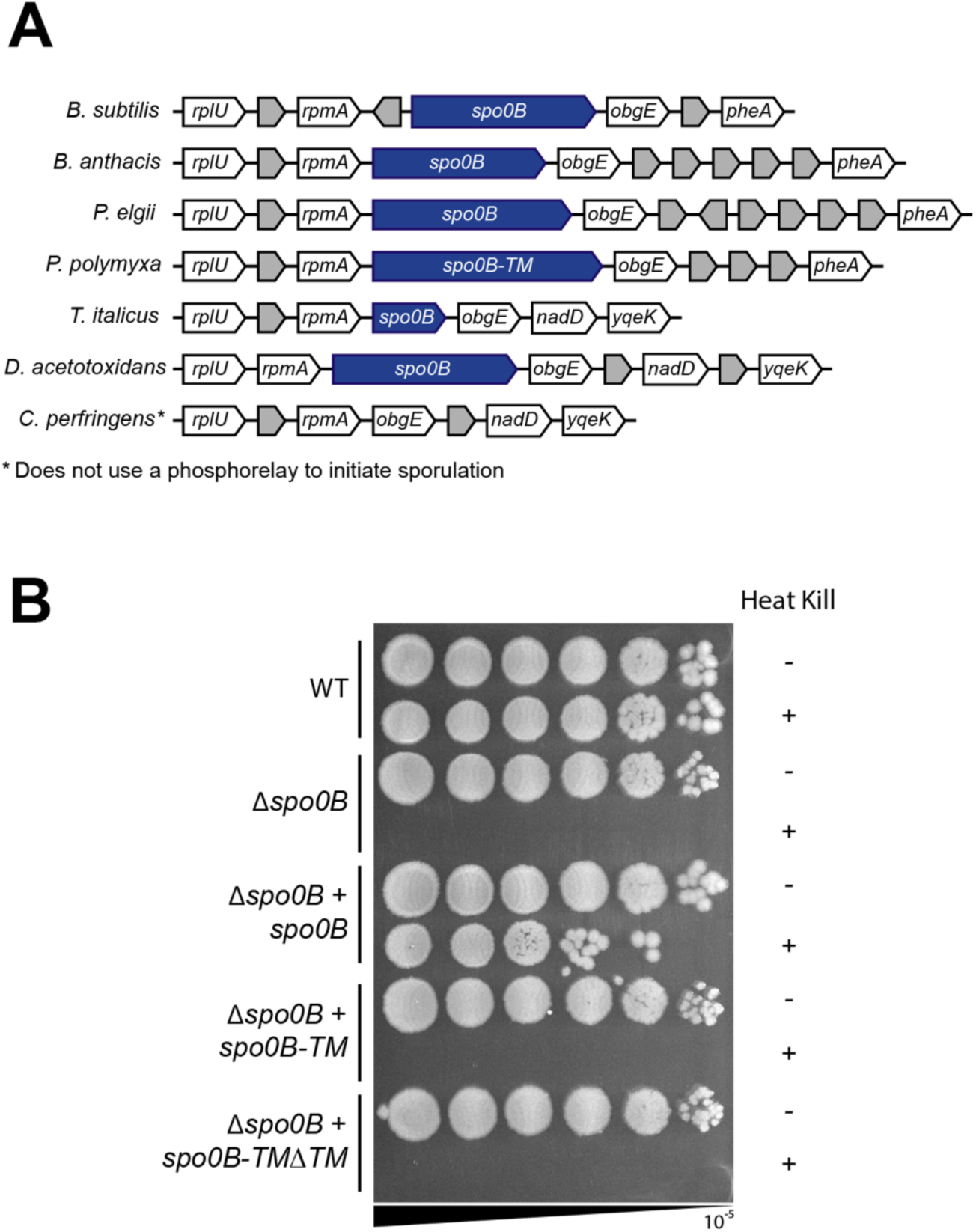
Spo0B-TM from *P. polymyxa* does not rescue the sporulation defect of a *spo0B* deletion in *B. subtilis*. **(A)** Synteny diagram of the Spo0B genomic region across *Bacillus subtilis, Bacillus anthracis, Paenibacillus elgii* (Spo0B), *Paenibacillus polymyxa* (Spo0B-TM), *Thermoanaerobacter italicus, Desulfofarcimen acetotoxidans,* and *Clostridium perfringens*. **(B)** Sporulation assay of *B. subtilis* strains. Serial dilutions of culture were plated before and after heating to 80°C to kill vegetative cells. Spot plate is representative of three biological replicates.

Because the genomic location and the phosphorylatable histidine domain of Spo0B-TM are conserved and Spo0B-TM can interact with other members of the *P. polymyxa* phosphorelay, we hypothesized that Spo0B-TM may be functional in related spore-formers. To test if *spo0B-TM* is functionally interchangeable with *spo0B* in *B. subtilis*, we expressed *P. polymyxa spo0B-TM* under the control of the native *spo0B* promoter and determined whether it could complement the *Δspo0B* mutant of *B. subtilis*. Spo0B is essential for sporulation in *B. subtilis*, and, as expected, the *Δspo0B* mutant failed to produce spores (Fig 5B). Complementation with *spo0B* under its native promoter restored sporulation efficiency to nearly wild-type levels (Fig 5B). The incomplete complementation may be because deleting *spo0B* has a polar effect on the downstream gene *obg*, a gene that is essential for both growth and sporulation (66, 67). Complementation with *spo0B-TM*, regardless of the presence or absence of its transmembrane domain, did not rescue the sporulation defect of the *Δspo0B* mutant, as no heat-resistant spores were detected in either strain (Fig 5B). Therefore, *spo0B-TM* and *spo0B-TMΔTM* from *P. polymyxa* are not functionally interchangeable with *spo0B* in *B. subtilis*, despite the highly conserved region surrounding the phosphorylatable histidine.

### Spo0B exhibits the lowest sequence identity of members of the phosphorelay

Since neither Spo0B-TM nor Spo0B-TMΔTM from *Paenibacillus* could rescue sporulation in a *Δspo0B* mutant of *B. subtilis*, we next examined the sequence conservation of Spo0B proteins across Bacilli. Despite a high degree of structural similarity predicted by AlphaFold (Fig 3B), *B. subtilis* and *P. polymyxa* Spo0B share only 19% protein sequence identity (Fig 6A). In contrast, Spo0F and Spo0A from these species share 70% and 67% protein sequence identity, respectively. Consistent with the low sequence identity of Spo0B, a search using BLASTN identified no significant Spo0B homologs in *Paenibacillus*. However, a search with BLASTP yielded hits for Spo0B in 1450/1460 genomes. Using HMMER, we were able to identify Spo0B in all but one of *Paenibacillus* genomes when we used the protein sequences of Spo0B from *P. polymyxa* and *P. validus* as the query.

**Figure 6.**
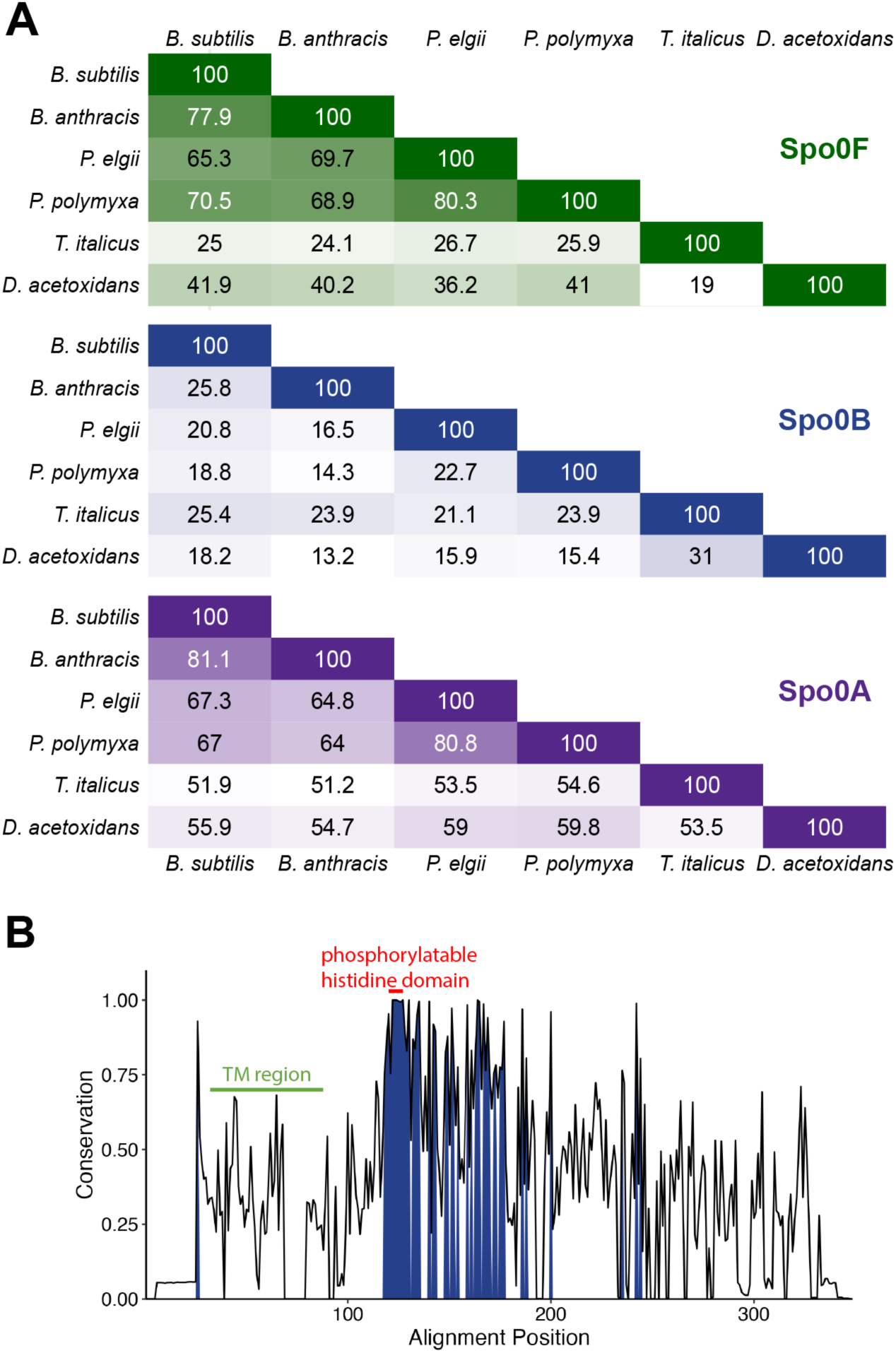
Spo0B protein sequences are not well-conserved. **(A)** Pairwise percent identity matrix of Spo0F, Sp00B, and Spo0A orthologs, calculated using Clustal Omega. **(B)** Protein sequence conservation plot of Spo0B-TM and Spo0B from *Paenibacillus* (n = 1437). Residues shared by ≥75% of sequences are shaded dark blue. Features are labeled with respect to the Spo0B-TM sequence from the reference genome of *P. polymyxa*.

The sequence of Spo0B is known to be poorly conserved among *Bacillus* species (68, 69), and we determined this to be true among *Paenibacillus* species as well (Fig 6B). In an alignment of 1437 Spo0B-TM and Spo0B sequences (length = 246±12 amino acids), six residues were 100% conserved (Fig 6B). Five of these residues are part of the highly conserved region around the phosphorylatable histidine residue. Only 46 residues were conserved in >75% of sequences (Fig 6B). The sequence of the transmembrane region was not strongly conserved, although its hydrophobic properties are maintained, suggesting that the importance of this domain is membrane localization. Altogether, these data suggest that the phosphorylatable histidine domain is the most fixed feature of this protein, while the remaining sequence space can evolve more freely.

## Discussion

Firmicutes diversified from other phyla very early in evolutionary history (70) and endospore formation is likely an ancestral trait of this phylum. Thus, the ability to form spores encompasses diverse bacteria that inhabit a broad range of environments. Sporulation genes have largely been inherited vertically, with approximately 120 genes shared across endospore formers within Firmicutes (23). Although previous studies included single representatives of *Paenibacillus*, we detected most of these genes in our expanded survey, which provides further support for their universal conservation. Core genes are involved in all stages of sporulation from initiation to germination, indicating that the fundamental process of sporulation is shared in most Firmicutes. Nevertheless, even within this core sporulation genome there is unexpected diversity that likely reflects differences in environmental contexts and signals triggering the decision to sporulate.

The ancestral sporulation initiation pathway is hypothesized to have been a phosphorelay (25), in which the phosphotransferases Spo0F and Spo0B aid in the transfer of the phosphoryl group from a histidine kinase to Spo0A (Fig 1). Spo0B, a cytoplasmic dimeric histidine transferase, may have evolved from a histidine kinase that lost its N-terminal signal detection domain and C-terminal ATP-binding domain (71, 72). Spo0B orthologs tend to vary greatly, even between species in the same genus (58, 68, 69) (Fig 3C, Fig 6). Park et al. noted that Spo0B in *P. polymyxa* has an extended N-terminal transmembrane region that is absent from Spo0B in *B. subtilis* (58). They found this unique transmembrane region in Spo0B sequences in a total of six *Paenibacillus* genomes. We identified Spo0B in all *Paenibacillus* genomes we surveyed (n = 1460), and of these, 92% encode Spo0B with an N-terminal transmembrane region (Spo0B-TM) (Fig 2). This transmembrane region was likely present in the ancestral *Paenibacillus* Spo0B and subsequently lost in certain lineages.

What is the importance of the transmembrane domain of Spo0B in *Paenibacillus*? Using a BACTH assay, we found that deleting the transmembrane domain eliminated detectable interaction between Spo0B and both Spo0A and Spo0F of *P. polymyxa* (Fig 4B). It is not immediately obvious how deleting this domain would disrupt interaction with either phosphorelay partner, but in *B. subtilis*, Spo0B forms dimers through the interaction of the N-terminal helical domains. Dimerization is essential to form the four-helix bundle interaction site necessary for phosphoryl transfer (73). Therefore, the additional helices supplied by the transmembrane domain could help to stabilize this formation and enable detection by BACTH. Species from *Paenibacillus* lineages that encode Spo0B without the transmembrane region have been experimentally shown to sporulate (74–76), which suggests that Spo0B lacking the transmembrane domain is functional in these lineages. Nevertheless, the high conservation of the transmembrane domain suggests it imparts a strong selective advantage in this genus. Future studies are needed to determine why localization of Spo0B to the membrane is important for Paenibacillus and whether this localization has a role in regulation of the phosphorelay or serves a sensory purpose.

Amongst members of the sporulation initiation phosphorelay we found that Spo0B exhibits the most sequence diversity (Fig 6A). The phosphorelay is an expanded version of a typical two-component system, and the interaction surface residues of the proteins in these systems resist evolutionary change. For example, 20 out of 21 interaction residues of Spo0F in *B. subtilis, Bacillus halodurans, B. anthracis* are identical, while only 50% of the remaining residues are identical (77). Accordingly, the most well-conserved region of Spo0B is the phosphorylatable histidine residue region (Fig 3B). Spo0B orthologs have a much lower sequence conservation, and in some instances, have even acquired additional function. For example, Spo0B in *B. anthracis* has pleiotropic functions, including phosphotransferase, autophosphorylation, and ATPase activity as a result of having acquired ATP binding and hydrolysis domains (69). The physiological consequence of gaining ATPase function has not been determined, but it has been proposed that this expanded functionality could be important for vegetative growth or even pathogenesis.

Our study highlights the importance of studying sporulation proteins in non-model organisms, since variations can be identified even in highly conserved sporulation genes.

*Paenibacillus* species inhabit a broad range of environments and cause a broad range of industrial impacts, from milk spoilage to bee-hive collapse. And yet, the majority of what we know about endospore formation comes from *B. subtilis*, mainly due to its genetic tractability. The growing number of *Paenibacillus* genomes available will greatly facilitate future surveys of this kind. Moreover, the wealth of environmental isolates that can be obtained from agricultural sources makes *Paenibacillus* an attractive and under-appreciated model of sporulation that may have important industrial applications.

## Data Availability Statement

Genome and protein accessions can be found in Tables S1 and S3. Scripts for data acquisition and analyses are available on GitHub at https://github.com/isabella-n-lin/paenibacillus_sporulation.

## Materials and Methods

### Strains and Media

*B. subtilis* strains were derived from 168 *trpC2* (78) and grown in Lysogeny Broth (LB) at 37°C with aeration. Genomic DNA from the *Δspo0B*::*kan* strain from the BKK collection (79) was used to create the Δ*spo0B* strains by transformation into 168 *trpC2*. Plasmids used can be found in Table 1. Antibiotics were used at final concentrations of 20 µg/mL kanamycin and 100 µg/mL spectinomycin. *Paenibacillus polymyxa* ATCC 842 was obtained from the American Type Culture Collection and grown in Brain Heart Infusion Broth at 30°C with aeration.

**Table 1.**
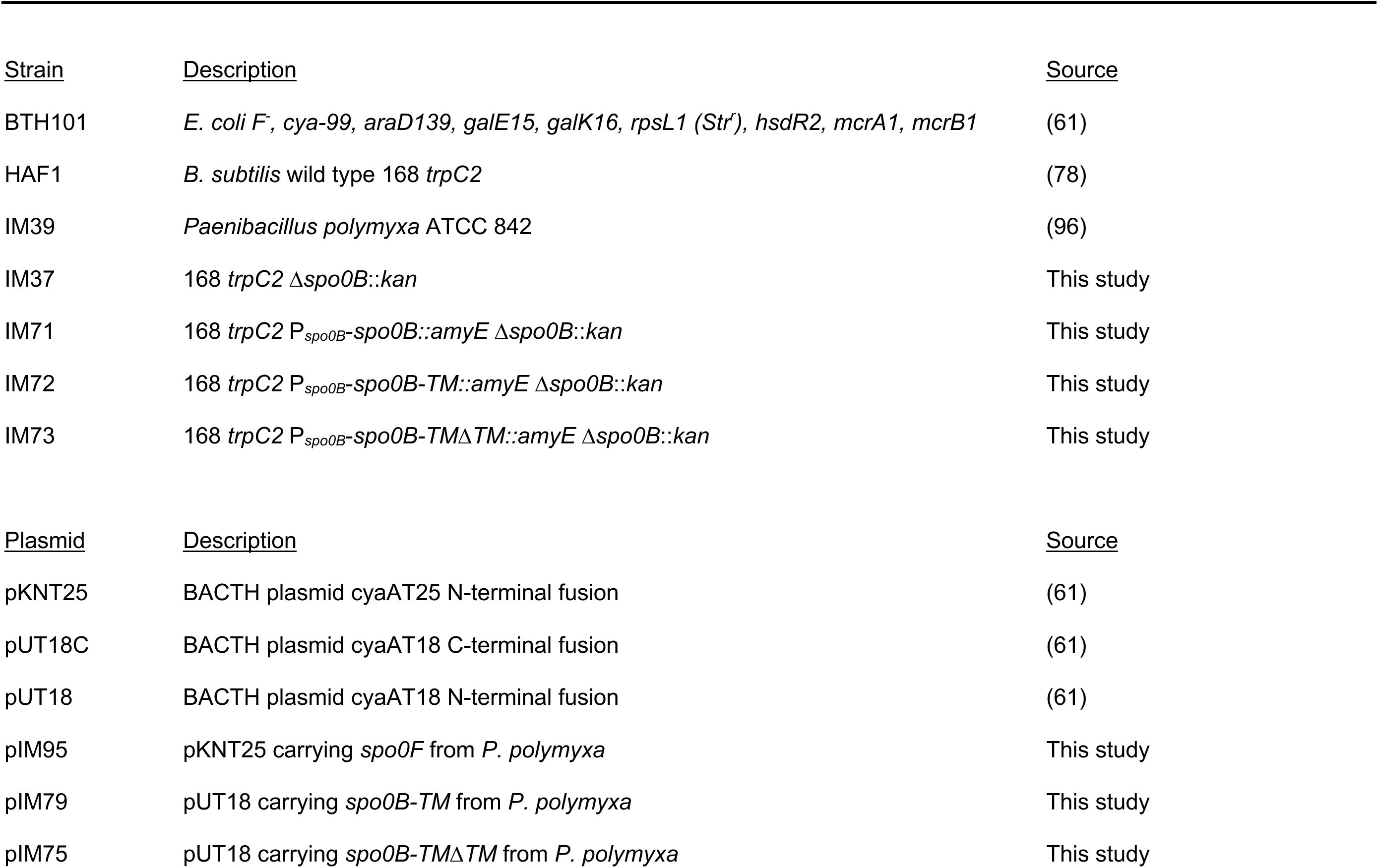
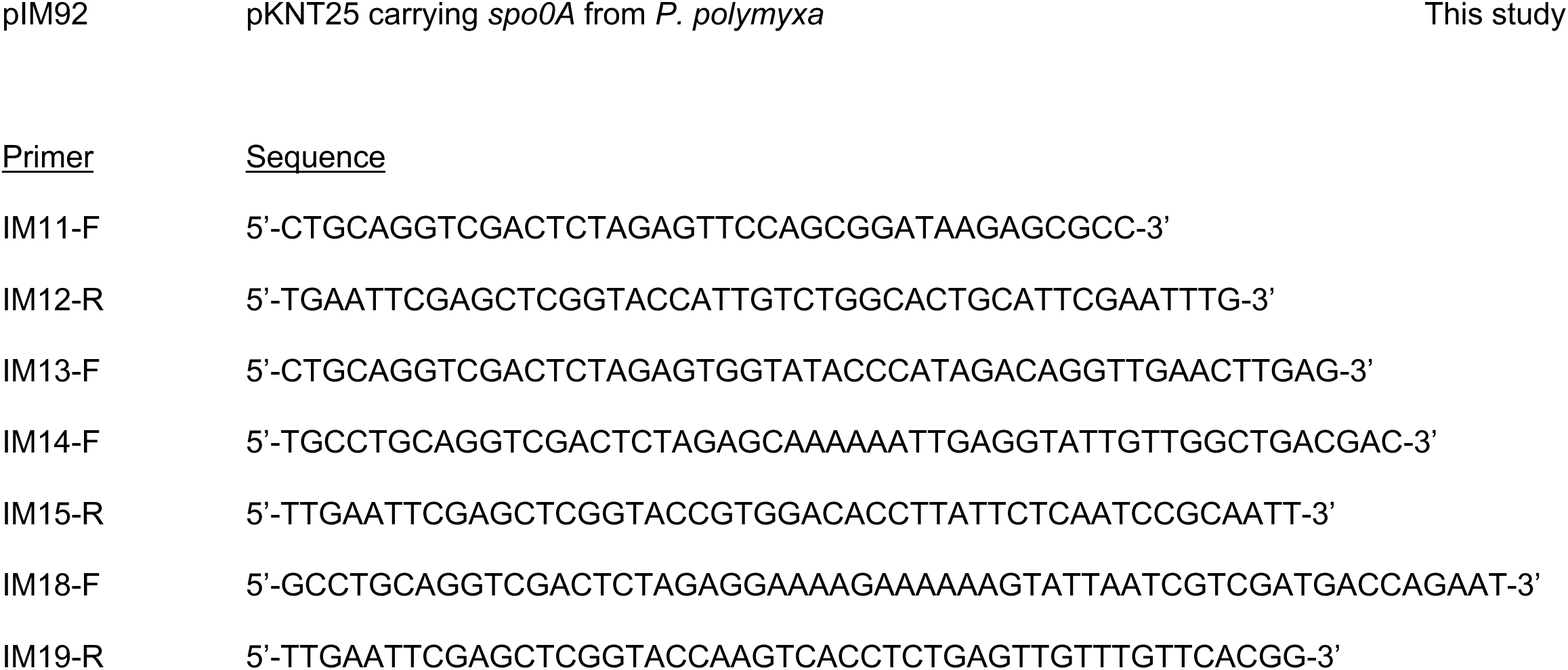
Strains, plasmids, and primers.

### Paenibacillus sporulation pangenome

All available *Paenibacillus* genome assemblies as of 21 April 2025 were downloaded from the NCBI Reference Sequence (RefSeq) collection as annotated by the NCBI Prokaryotic Genome Annotation Pipeline (80). Genomes with CheckM (81) contamination greater than 5% or completeness less than 95% were removed. Any genomes suppressed by NCBI by 28 July 2025 were removed, resulting in 1460 genomes. Genome assembly numbers can be found in Table S1. The phylogenetic tree was made with GToTree v1.8.14 (82), using the prepackaged single-copy gene set for bacteria (74 target genes). The tree was visualized using the ggtree v3.10.1 package (83) and midpoint rooted using the phangorn v2.12.1 package (84). A comprehensive list of sporulation genes was compiled from previous work (21, 23) and SubtiWiki (55) (Table S2). *B. subtilis* sporulation protein sequences were obtained from SubtiWiki (55). Sporulation genes were detected in *Paenibacillus* genomes using BLASTP v2.16.0 (85). HMMER v3.4 (86) was used to search for Spo0B across *Paenibacillus*, and DeepTMHMM v1.0.42 (87) was used to predict transmembrane regions. The protein accessions from *P. polymyxa* and *P. validus* for the HMMER query can be found in Table S3.

### Spo0F, Spo0B, and Spo0A sequence analysis and structure modeling

The Kyte-Doolittle hydropathy plot was created using ProtScale on the ExPASy server (88) using the Spo0B sequence WP_019688407.1. AlphaFold models were created using AlphaFold 3 (89) and visualized using ChimeraX v1.9 (90). Protein sequences of Spo0F, Spo0B, and Spo0A were obtained from NCBI (91) and the accession numbers can be found in Table S3. Percent identity was calculated using Clustal Omega v1.2.4 (92). The alignment in Figure 3C was made using Clustal Omega v1.2.4 (92) and visualized using the pyMSAviz v0.5.0 package (93). The conservation plot was created using the bio3d v.2.4-5 package (94). Spo0B sequences for the conservation plot were aligned using MAFFT v7.520 (95), using a gap opening penalty of 3 and gap extension penalty of 0.3.

### Bacterial two-hybrid system

To investigate the protein interactions of *spo0B-TM* from *P. polymyxa*, we used the commercially available Bacterial Adenylate Cyclase Two-Hybrid (BACTH) System Kit from Euromedex. Strains were derived from the non-reverting adenylate cyclase deficient *E. coli* strain BTH101. Plasmids used can be found in Table 1. Plasmid gene inserts were amplified from *P. polymyxa* ATCC 842. Antibiotics were used at final concentrations of 50 µg/mL kanamycin and 100 µg/mL ampicillin.

For the β-galactosidase assay, strains were grown in LB supplemented with 1% glucose, ampicillin, and kanamycin for 6 hours at 30°C. The cultures were used to inoculate LB supplemented with IPTG (0.5 mM), ampicillin, and kanamycin and grown for 18 hours at 30°C. The cells were lysed using lysis buffer (100 mL Z-buffer (0.06M Na_2_HPO_4_, 0.04M NaH_2_PO_4_-H_2_O, 0.01M KCl, 0.001M MgSO_4_-7H_2_O), 270 uL β-mercaptoethanol, 50 uL 10% sodium dodecyl sulfate) and 20uL chloroform and incubated at 30°C. The reaction was started upon the addition of o-nitrophenyl-β-D-galactopyranoside (4 mg/mL in Z-buffer) to the lysed cells. The reaction was stopped with 1M Na_2_CO_3_ when a pale-yellow color developed. Absorbances at 600, 420, and 550 nm were measured with a BioTek Synergy H1 microplate reader, Gen 5 3.11, for cell density, o-nitrophenol production, and background, respectively. β-galactosidase activity was measured in Miller units. *P*-values were calculated with the R stats v4.3.2 package using a one-way ANOVA followed by Tukey’s test. Experiments were performed using nine biological replicates.

### Sporulation assays

Sporulation was induced by nutrient exhaustion in Difco sporulation medium (DSM). Strains were grown in DSM on a roller drum at 37°C for 3 hours, then normalized to OD600 of 0.05 and incubated at 37°C, shaking at 220 rpm for 24 hours. The cultures were serially diluted in Tbase + 1mM MgSO_4_ and plated on LB agar plates before and after heating for 20 minutes at 80°C.

## Acknowledgements

HAF, INL, and CRP were supported by NIH R35GM147049. CRP was supported by a Graduate Research Fellowship from the National Science Foundation. This work was supported by the USDA National Institute of Food and Agriculture, Hatch project number 7005400.

